# Chromatin rewiring mediates programmed evolvability via aneuploidy

**DOI:** 10.1101/407841

**Authors:** Cedric A. Brimacombe, Jordan E. Burke, Jahan-Yar Parsa, Jessica N. Witchley, Laura S. Burrack, Hiten D. Madhani, Suzanne M. Noble

## Abstract

Eukaryotes have evolved elaborate mechanisms to ensure that chromosomes segregate with high fidelity during mitosis and meiosis^1^, and yet specific aneuploidies can be adaptive during environmental stress^2,3^. Here, we identify a chromatin-based system for inducible aneuploidy in a human pathogen. *Candida albicans* utilizes chromosome missegregation to acquire resistance to antifungal drugs^4,5^ and for ploidy reduction after mating^6^. We discovered that the ancestor of *C. albicans* and two related pathogens evolved a variant of histone H2A that lacks the conserved phosphorylation site for Bub1 kinase^7^, a key regulator of chromosome segregation^1^. Expression of this variant controls the rates of aneuploidy and antibiotic resistance in this species. Moreover, CENP-A/Cse4, the histone H3 that specifies centromeres, is depleted from tetraploid mating products and virtually eliminated from cells exposed to aneuploidy-promoting cues. Thus, changes in chromatin regulation can confer the capacity for rapid evolution in eukaryotes.

---

Aneuploidy is increasingly recognized as an important form of natural variation in eukaryotes. Chromosome segregation during mitosis and meiosis is generally accurate, but environmental stress has been associated with chromosome instability in multiple species^8,9^. Here, we investigate the molecular basis for stress- and ploidy-associated genome instability in the diploid (2n) yeast *Candida albicans.* This organism is the most common agent of fungal disease in humans, but primarily exists as a ubiquitous component of mammalian gut, skin, and genitourinary microbiota^10,11^. The ability of *C. albicans* to thrive as both a commensal and pathogen in diverse host niches has been attributed to its ability to transition among multiple specialized cell types^12^, as well as its remarkable genomic plasticity and tolerance of aneuploidy^9^. Aneuploidy is common in *C. albicans*, and is often a response to toxic stress^13^. For example, strains that exhibit resistance to the antifungal drug, fluconazole, frequently carry a duplication of the left arm of chromosome 5^4,5,14^, thereby increasing the copy number of the drug target as well as drug efflux pumps. The mechanisms of chromosome loss and aneuploidy generation in *C. albicans* are not well established; however, as described below, there is evidence supporting the involvement of tetraploid (4n) intermediates.

*C. albicans* tetraploid strains may be generated experimentally by exposure to fluconazole^15^ or by mating, wherein diploid *MTL***a** and *MTL*α cells mate to form tetraploid (4n) *MTL***a**/α progeny^16,17^. Tetraploid strains generally return to the diploid state either gradually, under most *in vitro* conditions, or abruptly, on certain types of media, by a process of random chromosome loss^6,18^. This unusual “parasexual cycle” is shared by *C. albicans* and the related pathogenic species *Candida tropicalis*^19^ and *Candida dubliniensis*^20^. In other ascomycetous fungi such as *Saccharomyces cerevisiae,* haploid **a** or a cells mate to produce diploid **a**/a progeny (reviewed in ^21^), which subsequently undergo meiosis to return to the haploid state. The mechanisms controlling random chromosome loss in *C. albicans* have remained elusive.

From yeast to humans, chromosome segregation is tightly regulated by a network of conserved checkpoint proteins. Regulated processes include the correction of erroneous microtubule attachment to sister chromatids and protection of centromeric cohesion during mitosis^1^. A central regulator is Bub1 kinase, which phosphorylates histone H2A on a highly conserved serine/threonine at position 121 (S/T121; **Fig 1A**). This mark subsequently recruits shugoshins^7,22^, which themselves recruit protein phosphatase 2A (PP2A) and the chromosomal passenger complex (CPC), containing Aurora B kinase. PP2A protects centromeric cohesin from degradation and promotes bi-orientation of sister chromatids prior to segregation^23^, whereas Aurora B ensures correct microtubule-kinetochore attachment prior to anaphase.

**Figure 1.**
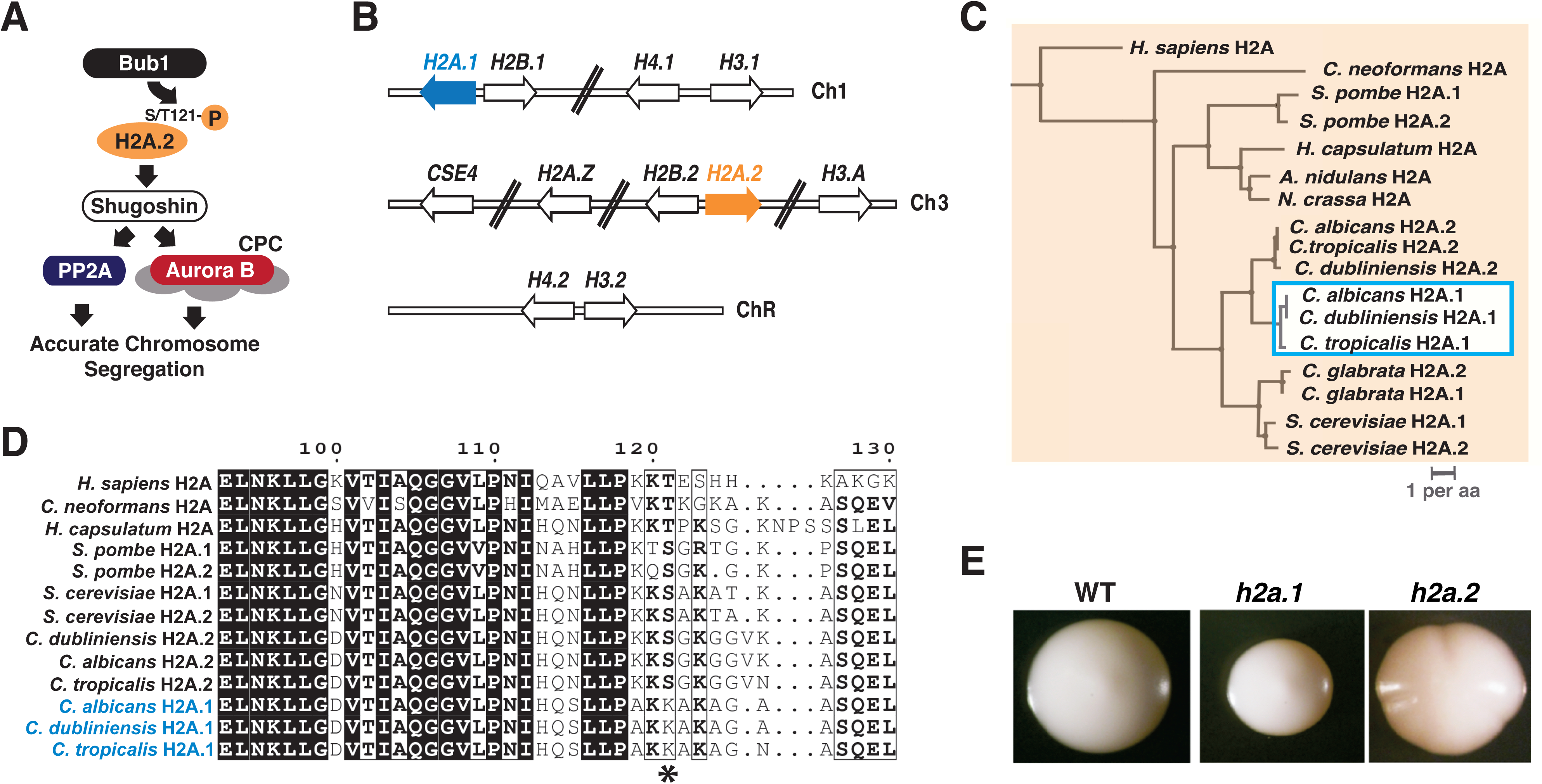
*C. albicans* encodes two alleles of Histone H2A that differ in C-terminal amino acid sequence. A) Schematic of Bub1-H2A.1-Sgo1 interactions in other eukaryotes. *C. albicans* retains orthologs of both Bub1 (orf19.2678) and Sgo1 (orf19.3550). B) *C. albicans* histone genes are distributed over three chromosomes. *H2A.1* (orf19.1051), *H2B.1* (orf19.1052), *H4.1* (orf19.1061), and *H3.1* (orf19.1059) occur on Chromosome 1, *CSE4* (orf19.6163), *H2A.Z* (orf19.327), *H2B.2* (orf19.6924), *H2A.2* (orf19.6925), and *H3.A* (orf19.6791) on Chromosome 3, and *H4.2* (orf19.1854) and *H3.2* (orf19.1853) are on Chromosome R, which also contains rRNA genes. C*) C. albicans, C. dubliniensis*, and *C. tropicalis* evolved divergent alleles of H2A. Among these three yeasts, *H2A.1* (orange shading) and *H2A.2* (blue shading) are more similar to their orthologs in the two other species than to the H2A paralog in their own species. Scale bar designates substitutions per amino acid. D) Amino acid differences between the H2A paralogs of *C. albicans, C. dubliniensis,* and *C. tropicalis* (not shown) localize to the C-terminus and include replacement of the highly conserved serine/threonine 121 residue with lysine (asterisk). E) Deletion of *H2A.2* but not *H2A.1* produces a colony sectoring phenotype. Wild-type and homozygous deletion mutants were propagated on YEPD medium and incubated at 30°C.

A search for homologs of Bub1, Sgo1, and H2A revealed that *C. albicans* contains single orthologs of Bub1 and Sgo1 (Figure 1A), as well as two homologous core H2A genes paired with divergently transcribed H2B genes, located on chromosomes 1 and 3 respectively (Figure 1B). We hereafter denote the H2A gene on chromosome 1, “*H2A.1*,” and that on chromosome 3, “*H2A.2*.” The primary amino acid sequences of H2A.1 and H2A.2 differ in the C-terminal region, most notably with the replacement of serine 121 with a lysine in H2A.1 but not in H2A.2 (Figure 1, B and C and Figure S1). A phylogeny of core H2A proteins revealed that the K121-associated allele (H2A.1) also occurs in *C. tropicalis* and *C. dubliniensis* (Figure 1C), two closely related yeast species that share a similar parasexual cycle with *C. albicans*. Because mutation of S/T121 results in chromosome instability in other organisms^7^, we hypothesized that deletion of *H2A.2* but not *H2A.1* would produce the same phenotype in *C. albicans*, and that the evolution of an H2A variant that is resistant to phosphorylation by Bub1 might offer a mechanistic explanation for the chromosome changes observed during the parasexual cycle and upon exposure to aneuploidy-inducing stresses.

To test this hypothesis, we generated knockouts of *H2A.1, H2A.2*, as well as the adjacent *H2B* genes as controls (**Figure 1D** and Fig S2). These experiments revealed that deletion of either *H2A.1* or *H2B.1* results in duplication of chromosome 3, on which the *H2A.2*-*H2B.2* locus is located; however the converse is not true when *H2A.2* or *H2B.2* is deleted (Figure S2, Table S1A). An analogous duplication of the *H2A-H2B* locus occurs in *S. cerevisiae* when one set of these loci is deleted, although in this case duplication occurs via the formation of a small circular episome rather than the entire chromosome^24^; a similar phenomenon occurs with reduction in histone H4 gene dosage in *C. albicans*^25^.

Because deletion of *H2A.1* or *H2B.1* results in duplication of chromosome 3, we used an alternative strategy to determine the phenotype of strains lacking these genes. As shown in Fig S3A, ‘ORF swapped’ strains were constructed in which both copies of the *H2A.1* open reading frame (ORF) are precisely replaced with the *H2A.2* ORF, and vice versa. The resultant strains contain either *H2A.1* or *H2A.2* at all four core H2A loci, and are thus deletion mutants for the other histone gene. Neither ORF-swapped strain contains aneuploidies, and measurements of genomic copy number and mRNA levels confirmed correct ORF replacement and transcription in the ORF swapped strains (Figure S3, A, B, and C). For clarity, we hereafter refer to these strains based on the copy number of each histone gene with the order [#*H2A.1*, #*H2A.2*], followed by ploidy; for example, a strain with four copies of *H2A.1* is denoted [4,0] diploid (see example in Figure S3A).

We tested the hypothesis that *H2A.2* but not *H2A.1* is required for chromosome stability by comparing the phenotypes of strains lacking either histone gene. On standard growth medium, the strain lacking *H2A.1* ([0,4] diploid) is indistinguishable from WT; however, the strain lacking *H2A.2* ([4,0] diploid) exhibits sectoring in roughly 25% of colonies (Figure 2, A and B). Colony sectoring is often symptomatic of chromosome instability, with defects in colony margins reflecting clones of inviable cells. Consistent with this interpretation, deletion mutants affecting the upstream (Bub1) and downstream (Sgo1) spindle assembly checkpoint components also exhibit sectoring phenotypes (Figure 2, A and B). The *sgo1* mutant sectors to the same degree as the [4,0] strain, whereas *bub1* has an even stronger phenotype, likely reflecting additional checkpoint functions of Bub1^1,7^. We queried the role of the S121 residue of H2A.2 on the phenotype by testing the ability of wild-type *H2A.2* vs. a mutant differing by a S121A substitution to suppress colony sectoring. When restored to the genome of the [2,0] strain, wild-type *H2A.2* reduces sectoring to 2% of colonies, but the S121A variant has no effect (Figure 2, A and B). These results suggest that increased colony sectoring is linked to H2A alleles that are not substrates for Bub1 kinase.

**Figure 2.**
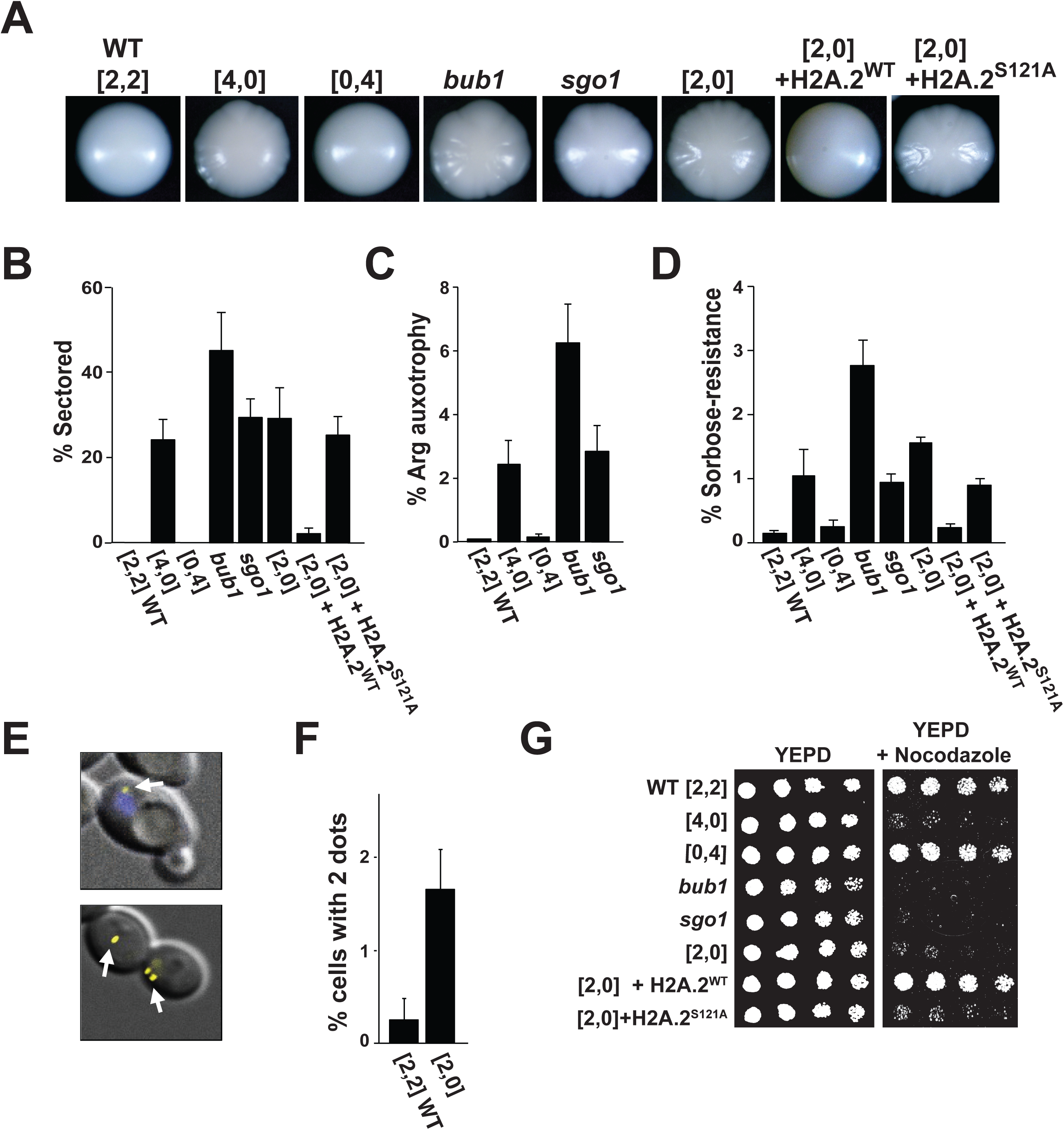
A *C. albicans* mutant lacking *H2A.2* phenocopies spindle checkpoint mutants. A and B) Deletion of *H2A.2, BUB1, SGO1*, and replacement of *H2A.2* with a *H2A.2* S121A mutation result in similar colony sectoring phenotypes. Colony morphology (A) and % colony sectoring (B) are shown for wild-type *C. albicans*, the [4,0] strain that lacks *H2A.2*, the [0,4] strain that lacks *H2A.1*, knockouts of the spindle checkpoint genes, *BUB1* and *SGO1*, and the [2,0], and [2,0]+*H2A.2*S121A mutants. C, D, E, and F) Mutants lacking *H2A.2, BUB1*, and *SGO1* display increased aneuploidy. C) Percent of colonies that have lost the ability to grow on arginine-deficient medium, relative to growth on YEPD. Strains were plated on arginine-deficient medium or YEPD after 4 days on pre-sporulation medium. Because these *ARG4/*D strains contain the *ARG4* gene on only one copy of chromosome 7, loss of that chromosome renders cells auxotrophic for arginine. D) Percent of colonies that grow on sorbose-containing medium, relative to growth on YEPD. Because cells with both copies of chromosome 5 are killed by sorbose, this medium selects for cells that have lost one copy of the chromosome. E) Top, Superimposed fluorescence and phase microscopy of QMY85, which contains tandem copies of the tet operator on Chromosome 5, as well as the tet repressor fused to YFP. A yellow spot marks one of two copies of Chromosome 5 (white arrow) and the nucleus is stained with DAPI (blue arrow). Bottom, example of neighboring cells with 1 or 2 spots. F) % of cells with two spots, counting un-budded cells only. G) Sensitivity to the microtubule poison Nocodazole (50 µM) shown by serial dilution.

We next assessed chromosome stability in these strains by introducing heterozygosity for the auxotrophic marker *ARG*4 (located on chromosome 7), followed by testing for marker loss over time. Compared to wild type or the [0,4] mutant, strains that lack H2A.2, Bub1, or Sgo1 exhibit highly elevated rates of marker loss that correlate with their colony sectoring phenotypes (Figure 2C). Additionally, we evaluated growth on sorbose-containing medium, which selects for cells that have lost a single copy of chromosome 5, with a similar pattern of results (Figure 2D); in this assay, restoration of one allele of wild-type *H2A.2* to the [2,0] strain reduced the number of colonies that grow on sorbose, whereas the S121A-containing variant again had no effect (Figure 2D). In a reciprocal experiment, we tested for chromosome gain events using a [2,0] *h2a.2* strain in which one copy of chromosome 5 can be visualized as a fluorescent dot, produced by the binding of a tetR-YFP fusion protein to >100 tandem copies of the *tet* operator (Figure 2E). As shown in Figure 2F, the frequency of un-budded cells that contain two nuclear dots is approximately six times higher in the *h2a.2* ([2,0]) mutant than in wild-type *C. albicans*. Finally, all strains that lack wild-type H2A.2—including the *h2a.2* strain that contains the S121A variant—as well as *bub1* and *sgo1* are hypersensitive to the microtubule-destabilizing drug nocodazole (Figure 2G), as would be expected for mutants affecting the spindle assembly complex^7^. The nocodazole-hypersensitive strains are also hypersensitive to ultraviolet radiation (Figure S4), consistent with an additional role for *C. albicans* H2A S121 in DNA repair as has been observed in other yeasts^7^. These results support our strong expectation, based on studies of other organisms, that H2A.2 but not H2A.1 acts as a substrate of the Bub1 kinase to recruit shugoshin to centromeric chromatin and thereby to ensure faithful chromosome segregation (Figure 1A). However, they raise the question of why *C. albicans* and closely related yeasts would maintain a gene encoding a defective histone H2A.

Based on the observations that 1) reduced H2A.2 levels result in chromosome instability, 2) *C. albicans* tetraploid cells exhibit an enhanced rate of random chromosome loss (as depicted in Figure 3A), and 3) only the three closely-related yeast species that undergo parasexual mating contain the H2A.1 variant (Figure 1, B and C), we posited that modulation of H2A.2 abundance could mediate ploidy reduction in tetraploid cells. To test this hypothesis, we compared the chromosome stability of wild-type tetraploid cells ([4,4] tetraploid) to that of tetraploids containing only H2A.1 ([8,0] tetraploid) or only H2A.2 ([0,8] tetraploid); ploidy was verified by flow cytometry (Figure S5, A and B). Viability and genomic marker loss were assayed on pre-sporulation (pre-spo) medium, which strongly promotes ploidy reduction in tetraploid cells^6^. Note that concerted chromosome loss is associated with reduced cell viability^6^. We observed that, after four days of propagation on pre-spo medium, the [4,4] tetraploid and [0,8] tetraploid strains exhibited ∼40 to 50% viability, whereas the viability of the [8,0] tetraploid strain was more drastically reduced to only ∼4% (Figure 3B); by comparison, there were no viability differences among these strains when propagated on standard YEPD medium (Figure 3B). The decreased viability of the [8,0] tetraploid strain that lacks canonical H2A.2 was associated with a significantly elevated rate of chromosomal marker loss compared to the [4,4] wild-type tetraploid strain. By contrast, the [0,8] tetraploid strain that lacks variant H2A.2 exhibited a significantly reduced rate of marker loss (Figure 3C). In summary, with respect to chromosome stability, tetraploids containing only H2A.2 are stabilized compared to wild-type tetraploids, and tetraploids with only H2A.1 are de-stabilized; colony sectoring phenotypes also correlated with the increased instability and death observed in these strains (Figure S4C).

**Figure 3.**
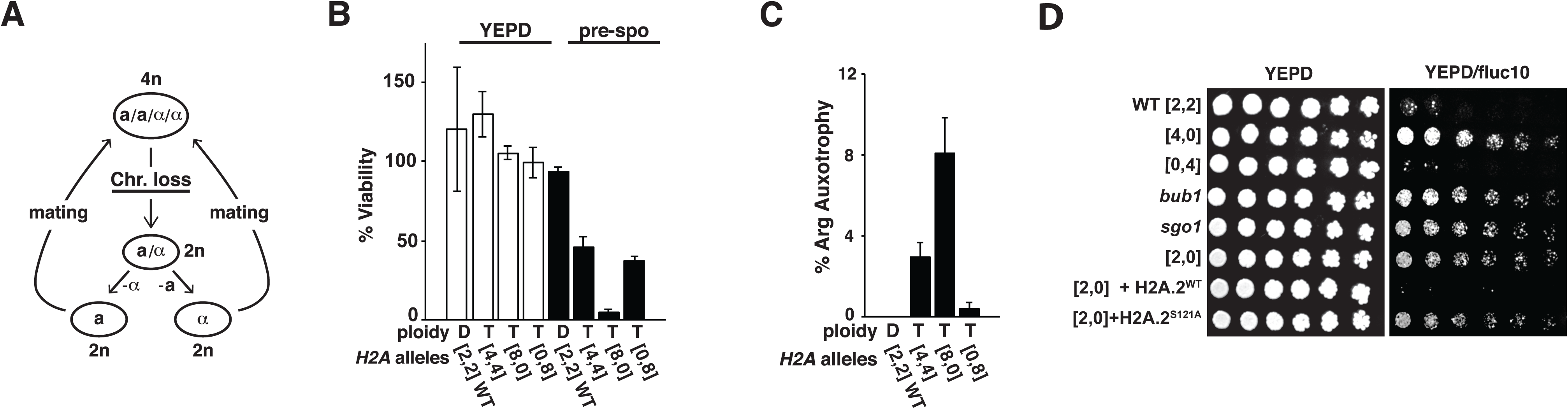
H2A.1 and H2A.2 play opposing roles in the chromosome stability of tetraploid cells and in the acquisition of resistance to fluconazole. A) Cartoon of the *C. albicans* parasexual cycle, which consists of mating between diploid **a** and a cells to form tetraploid **a**/a cells, followed by ploidy reduction by concerted chromosome loss rather than meiosis, as occurs in other eukaryotes. B) Viability of WT, tetraploid, or tetraploid cells containing only H2A.1 ([8,0] Tetraploid), or H2A.2 ([0,8] Tetraploid) on YEPD (left) vs. pre-sporulation medium (right). Plated cells were propagated for four days at 37°C, and survival was calculated as the ratio of colonies on each medium to the estimated number of plated cells determined using a hemocytometer. C) Chromosome stability of *ARG4/arg4*D WT, tetraploid, and derivative strains on pre-sporulation medium after four days, as measured by growth on SC(-Arg) medium, indicative of the loss of one copy of chromosome 7. D) Serial dilutions of *C. albicans* WT, [4,0] diploid, [0,4] diploid, *bub1, sgo1*, [2,0] diploid, and [2,1] diploid strains on YEPD or YEPD + fluconazole (10 µg/ml).

Because fluconazole resistance is known to be triggered by aneuploidy^15^, we additionally evaluated the development of fluconazole resistance in wild type ([2,2] diploid), [4,0] diploid, [0,4] diploid, *bub1, sgo1, h2a.2, H2A.2* add-back, and S121A strains. We observed that the [4,0] diploid, *bub1, sgo1, h2a.2,* and S121A strains developed fluconazole resistance at an elevated frequency compared to wild type, [0,4] diploid, or *H2A.2* add-back strains (Figure 3D).

We considered several potential mechanisms for the increased chromosome instability observed in tetraploid cells compared to diploid cells (Figure 3C), tetraploid cells propagated in pre-spo medium compared to YEPD (Figure 3B and ^6^), and diploid cells treated with fluconazole compared to untreated^15^. First, we asked whether the expression of the variant H2A.1 is upregulated relative to the canonical H2A.2 in cells that are susceptible to chromosome instability. However, RT-qPCR revealed no significant differences in *H2A.1* and *H2A.2* mRNA levels between any of these cell types or conditions (Figure 4A). Likewise, levels of an epitope-tagged H2A.2 fusion protein (H2A.2-FLAG) were similar in diploids, tetraploids, and diploid cells exposed to fluconazole (Figure 4B; note that epitope-tagged H2A.1 could not be obtained despite multiple attempts). Next, we asked whether the abundance of H2A.2 at centromeres and/or pericentromeric DNA might be reduced relative to non-centromeric regions in tetraploid cells. However, ChIP-Seq analysis of H2A.2-FLAG localization revealed only a slight, albeit statistically significant, reduction in tetraploid vs. diploid cells (p<0.05 for all pairwise comparison of replicates, Mann-Whitney U tests; Figure 4, C and D and Figures S6 and S7). In parallel, control ChIP-seq experiments performed with anti-H3 antibodies revealed depletion at centromeres in both cell types (Figure 3, C and D and Figures S6 and S7), as expected if canonical histone H3 is replaced by the centromere-specific histone H3, Cse4^26^ (CENP-A in other organisms^27^) in these regions.

**Figure 4.**
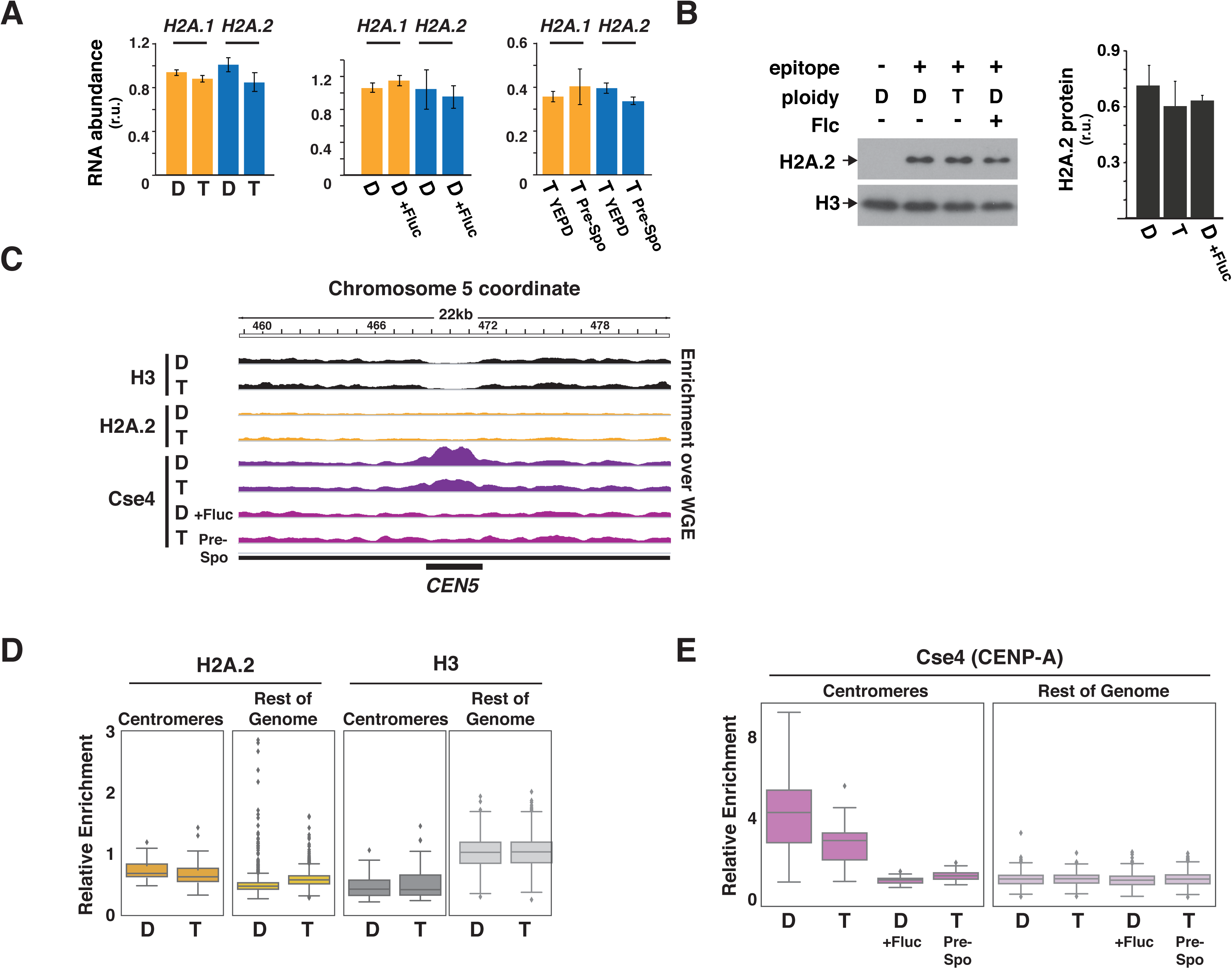
Impact of ploidy and environment on histone expression and deposition. A) RT-qPCR analysis of expression of *H2A.1* and *H2A.2* in the indicated genotypes and conditions. “D” represents a wild-type [2,2] Diploid strain and “T” a wild-type [4,4] Tetraploid strain. The average results for three biological replicates is presented after normalization to H3 mRNA, along with the standard deviation. R.u. stands for relative units. B) Immunoblot of levels of H2A.2-FLAG in the indicated genotypes and conditions. Quantification using a LI-COR imager is shown on the right. C) ChIP-seq analysis. Plots of bedgraphs of normalized read densities are shown for the indicated genotypes and conditions. Data represent the results of one replicate of each sample for *CEN5*. Data for all centromeres are shown in Fig. S6, and scatterplots of replicate data are shown in Fig. S7. D) Boxplots showing normalized ChIP-seq enrichments (relative to the whole-cell extract sample, see Materials and Methods) of nonoverlapping 5 kb tiles of the *C. albicans* genome separated into those overlapping with annotated centromeres vs. the remainder of the genome.

These data indicate that, while we have demonstrated that newly evolved H2A.1 facilitates chromosome gain and loss events in *C. albicans*, ploidy-associated differences in chromosome stability do not appear to result from differences in the relative expression or chromosome localization of H2A.1 versus H2A.2. We hypothesized that the presence H2A.1 may serve instead to sensitize cells to changes in other key centromere and kinetochore components. To investigate a potential role for Cse4/CENP-A, which forms the platform for assembly of eukaryotic kinetochores^27^, we performed ChIP-seq with polyclonal antibodies against this centromere-specific histone H3. Replicate analysis revealed that Cse4 is enriched at all seven non-repetitive centromeres (the repetitive R chromosome was excluded from analysis), as expected; however, the level of Cse4 was significantly reduced in tetraploid cells compared to diploids (p<1x10^-5^ for all four pairwise replicate comparisons; Mann-Whitney U tests; Figures 4C, 4E, S6, and S7). To determine whether Cse4/CENP-A deposition is also influenced by *in vitro* conditions associated with enhanced chromatin instability^6,15^, we repeated the analysis on extracts prepared from diploid cells exposed to 10 mg/ml fluconazole and tetraploid cells propagated in pre-spo medium. Under chromosome-destabilizing conditions, we observed a striking depletion of Cse4 at all centromeres (Figures 4C, 4E, S6, and S7; p<1x10^-15^ for all four pairwise replicate comparisons of diploid cells and p<1x10^-13^ for equivalent comparisons of tetraploid cells; Mann-Whitney U tests). These data indicate that at least two parallel pathways promote regulated chromosome instability in *C. albicans*: a newly evolved H2A lacking the Bub1 phosphorylation site sensitizes cells to small changes in centromere and kinetochore components, and a second mechanism modulates the centromeric accumulation of Cse4/CENP-A in response to changes in ploidy and the environment.

Chromosome instability is proposed to play key roles in *C. albicans* cell biology, including reduction of chromosome number after mating and adaptation to drug challenges. Although substantial evidence supports the ability of environmental perturbations to trigger chromosome instability in this organism ^6,13,15,28^, the underlying molecular mechanisms have remained obscure. We discovered that *C. albicans* and the related human pathogens *C. tropicalis* and *C. dubliniensis* have evolved a variant histone H2A protein, H2A.1, that is defective in chromosome segregation because of loss of an otherwise universally conserved phosphorylation site of the Bub1 kinase. We show that H2A.1 promotes ploidy modulation in the tetraploid products of mating and enhances the frequency with which fluconazole resistance occurs in a cell population. Moreover, we observed that deposition of the centromeric histone H3, Cse4/CENP-A, is regulated in this species by both ploidy and environment, suggesting that at least two histone-based mechanisms control chromosome stability. Thus, chromatin-based regulation of chromosome segregation enables parasexuality and the rapid acquisition of drug resistance in *C. albicans* (and, potentially, *C. dubliniensis* and *C. tropicalis*).

Variation is essential for evolution, and aneuploidy is a surprisingly common form of natural variation. For example, 19% of natural isolates of *S. cerevisiae* are aneuploid^29^, despite the fact that mitosis is a high-fidelity process in this species, at least under standard laboratory conditions. While aneuploidy is typically deleterious, specific aneuploidies have been shown to confer selective advantages to organisms ranging from fungi^2^ to cancer cells^3^ in times of stress. Given that many eukaryotes are subject to severe and unpredictable environmental fluctuations, the capacity for programmed variation through promiscuous chromosome segregation may be more general than currently appreciated.

## ACKNOWLEDGMENTS

We are grateful to Quinn Mitrovich and Sandy Johnson for their gift of the unpublished tetO strain, QMY85. This study was supported by NIH grants R01AI108992 to S.M.N and R01AI120464 to H.D.M. J.W. was supported by US National Institutes of Health grant T32AI060537. J.E.B. was supported by an American Cancer Society Postdoctoral Fellowship. H.D.M. is an Sr. Investigator of the Chan-Zuckerberg Biohub. S.M.N. is a Burroughs Wellcome Investigator of the Pathogenesis of Infectious Diseases and a Pew Scholar in the Biomedical Sciences.

## METHODS

### Media and growth conditions

Cells were propagated in yeast extract peptone dextrose (YEPD [2% bacto peptone, 1% yeast extract, 2% dextrose] or pre-sporulation medium [10% glucose, 0.8% yeast extract, 0.3% peptone] as previously described ^6^. For colony morphology analyses, cells were propagated at 30°C for 5 days on YEPD agar. For chromosome stability analyses of diploid strains, cells were propagated on pre-spo [10% glucose, 0.8% yeast extract, 0.3% peptone] agar medium at 37° C for four days; for chromosome loss assays in tetraploid strains, cells were propagated on pre-spo for four days at 37°C. For mRNA and protein analyses, cells were propagated in YEPD on a 30°C shaker running at 200 r.p.m to an OD of ∼1.0 before harvest for use in subsequent assays. For fluorescence microscopy analyses, cells were grown similarly to those prepared for mRNA analyses.

### Strain construction

A detailed description of strains, all of which are derivatives of the clinical isolate SC5314^30^, is provided in Table S2. *C. albicans* mutant and epitope-tagged strains were created as previously described.^31^ orf19 standard designations of genes newly named in this work include: *H2A.1* (orf19.1051), *H2B.1* (orf19.1052), *H2A.2* (orf19.6925), *H2B.2* (orf19.6924), *H3.1* (orf19.1059), *H4.1* (orf19.1061), *H3.2* (orf19.1853), *H4.2* (orf19.1854), *H3.A* (orf19.6791), *H2A.Z* (orf19.327).

#### ORF-swapping

All plasmids and primers used in strain construction and diagnosis are described in Table S3 and Table S4, respectively. H2A ORF-swapped strains were created using plasmids pSN379 and pSN380. pSN379 contains 300 bp upstream of the *H2A.1* ORF fused to *H2A.2* ORF (amplified from genomic DNA using primers SNO2811 and SNO2812), the *H2A.2* ORF (amplified from genomic DNA using SNO2313 and SNO2314), a *FLP-SAT* cassette ^32^ (amplified from pSFS2 ^32^ using SNO2815 and SNO2816), and 300 bp downstream of the *H2A.1* ORF (amplified from genomic DNA using SNO2017 and SNO2818), flanked on either side by PmeI restriction sites. pSN380 contains 300 bp upstream of the *H2A.2* ORF fused to the *H2A.1* ORF (using SNO2819 and SNO2820), the *H2A.1* ORF (using SNO2821 and SNO2822), the *FLP-SAT* cassette (using SNO2823 and SNO2824), and 300 bp downstream of the *H2A.2* ORF (using SNO2825 and SNO2826), flanked on either side by PmeI restriction sites. These constructs were assembled by homologous recombination in *S. cerevisiae* ^33^. Plasmids were linearized by digestion with PmeI and transformed into the WT strain SN152, with selection for nourseothricin-resistant colonies (NatR). Correct integration was verified by colony PCR targeting the 5’ and 3’ junctions using the primers SNO2859 and SNO2867, and SNO2862 and SNO2868 for integration of pSN379, and SNO2863 and SNO2867, and SNO2866 and SNO2868 for integration of pSN380. Cells were propagated in YEPD+2% maltose to permit excision of *FLP-SAT* cassette, with correct excision verified by PCR of the 5’ and 3’ junctions. Each Nat-sensitive heterozygous strain was then transformed a second time with the same linearized plasmid, again selecting for NatR colonies. Colony PCR was performed to confirm the absence each ORF, using SNO2735 and SNO2736 for *H2A.1*, and SNO2549 and SNO2550 for *H2A.2*. Colonies that were ORF negative for each opposite histone were validated for correct insertion by PCR across the 5’ and 3’ junctions. The *FLP-SAT* cassette was subsequently flipped out, with correct excision verified by junction PCR. The resultant strains contained four copies of *H2A.1* or *H2A.2*, with two copies in the native locus, and two copies in the other *H2A* locus, with FRT sites immediately 3’ of the stop codon of each gene. The absence of *H2A.1* native junctions was verified using SNO2661 and SNO2972 (5’), and SNO2662 and SNO2971 (3’). The absence of *H2A.2* native junctions was verified using SNO2547 and SNO2974 (5’) and SNO2603 and SNO2973 (3’).

#### Targeted Gene Disruption

Homozygous knockout mutants of *H2A.1, H2B.1, H2A.2, H2B.2, BUB1, SGO1, and MTL***a** were constructed by fusion PCR as previously described ^34^. For each strain, a brief description of primers used is as follows: *H2A.1* deletion constructs were created using the primers SNO2657-SNO2660, correct integration verified using SNO2661, SNO2662, and SNO265-268. Strains were verified as ORF negative using the primers SNO2735 and SNO2736. *H2A.2* deletion constructs were created using the primers SNO2543-SNO2546, correct integration verified using SNO2547, SNO2603, and SNO265-268. Strains were verified as ORF negative using the primers SNO2549 and SNO2550. *H2B.1* deletion constructs were created using the primers SNO2748-SNO2751, correct integration verified using SNO2752, SNO2753, and SNO265-268. Strains were verified as ORF negative using the primers SNO2725 and SNO2726. *H2B.2* deletion constructs were created using the primers SNO2665-SNO2668, correct integration verified using SNO2669, SNO2670, and SNO265-268. Strains were verified as ORF negative using the primers SNO2729 and SNO2730. *BUB1* deletion constructs were created using the primers SNO2999-SNO3002, correct integration verified using SNO3003, SNO3004, and SNO265-268. Strains were verified as ORF negative using the primers SNO3005 and SNO3006. *SGO1* deletion constructs were created using the primers SNO3086-SNO3089, correct integration verified using SNO3090, SNO3091, and SNO265-268. Strains were verified as ORF negative using the primers SNO3092 and SNO3093. *MTL***a** deletion constructs were created using the primers SNO2502-SNO2505. Strains were verified as ORF negative using the primers SNO2358 and SNO2359.

A homozygous knockout of *H2A.2* in the TetO-TetR-YFP strain was generated using the plasmid pSN383, with two successive rounds of transformation followed by excision of the *FLPSAT* cassette. pSN383 contains ∼300 bp upstream of the *H2A.2* ORF (amplified from genomic DNA using primers SNO3145-3146), the *FLP-SAT* cassette (amplified from pSFS2 using primers SNO3149-3150), and ∼300 bp downstream of the *H2A.2* ORF (amplified from genomic DNA using primers SNO3147-3148); both sides were flanked by PmeI restriction sites for linearization. Constructs were assembled by homologous recombination in *S. cerevisiae*, and sequenced. Linearized plasmid was transformed into *C. albicans* strain QMY85, and NatR colonies were selected; correct 5’ and 3’ integration events were confirmed using the primers SNO2543 and SNO1902, and SNO2546 and SNO1903. The FLP-SAT cassette was excised as described above, and a second round of transformation and *FLP-SAT* excision was performed. Strains were verified as ORF negative for *H2A.2* using the primers SNO2549-2550.

#### Complementation with H2A.2 and H2A.2 S2121A

The *H2A.2* gene addback construct was generated by precisely replacing the ORF of a previously used addback construct ^35^ with the *H2A.2* ORF, flanked by ∼1000 bp 5’ and 3’ of the gene, resulting in pSN381. Primers SNO3068 and SNO3071 were used to amplify the backbone of the plasmid, and SNO3068 and SNO3069 to amplify the inserted sequence. Products were then treated with DpnI, mixed in a 1:1 ratio, and transformed into *E. coli*, followed by sequencing of transformants to verify correct DNA sequence replacement. The S121A mutation was introduced by synthesizing a *H2A.2* gene that contains a TCA to GCA codon substitution at this position; the mutant ORF was subsequently swapped into the original addback construct by homologous recombination in *S. cerevisiae* ^33^, generating pSN382 using the same strategy as above, with correct replacement verified by sequencing. Each resultant plasmid contains several hundred bp of genomic sequence immediately upstream of the *LEU2* ORF, followed by the *H2A.2* (WT) or *H2A.2* (S121A) gene, *C.dubliniensis ARG4*, and several hundred base pairs of genomic sequence immediately downstream of the *LEU2* ORF, all of which is flanked by PmeI restriction sites, identical to previously described addback plasmids ^36^. Plasmids were linearized by digestion with PmeI, and transformed into the [2,0] diploid strain, with selection for Arg^+^ transformants. Presence of the ORF and the expected 5’ and 3’ junctions was verified by colony PCR using the primers SNO2549-2550 (ORF), SNO464+SNO2550 (5’), and SNO467+SNO2549 (3’).

#### Tetraploid mating products

[4,4], [8,0], and [0,8] tetraploids were generated in several steps. First, an allele of the Mating Type-Like Locus (*MTL*) was disrupted in [2,2], [0,2] and [2,0] diploid strains using plasmid pJD1 ^37^, which indiscriminately replaces either the *MTL***a** or *MTL*α locus with the *C.d.ARG4* gene. From these transformants, *MTL***a** cells were identified (i.e. disruptants of the *MTL*α allele), with correct deletion verified by colony PCR using the primers SNO2358-59 (*MTL*a) and SNO2360-61 (*MTL*α). In parallel, the *MTL***a** locus was deleted in the same starting strains using fusion PCR to replace the *MTL***a** locus with *C.d.HIS1* (or *C.m. LEU2* in the H2A.2-FLAG strain); correct transformants were identified by colony PCR. *MT*L**a**/D and *MTL*α/D strains were switched to the opaque phase and then mated together on YEPD agar medium for 8 hours. Mating products were selected on SD-arg-his or SD-arg-leu medium. Ploidy of the tetraploid mating products was verified by flow cytometry.

#### Epitope tagging of H2A.2

A strain containing a C-terminal fusion of *H2A.2* to the 6His/FLAG epitope was created using primers SNO2731-2732 and the pADH52 ^31^ template to amplify a *FLP-SAT-*marked PCR product containing the epitope and sequences flanking the *H2A.2* stop codon. The PCR products was transformed into SN152, and correct integration was verified by colony PCR using the primers SNO2549 + SN2288, and SNO2550 + SNO2289. The *FLP-SAT* cassette was then excised by growth in 2% maltose, and correct excision verified by PCR using the primers SNO2549 + SNO2603.

### Reverse transcription quantitative PCR analyses

RNA was extracted from log or stationary phase cultures grown in YEPD or Pre-spo medium, and extracted using a hot acid phenol method, as previously described ^17^. RNA was then treated with DNaseI (Ambion) for 1 hour at 37°C, followed by a single phenol-chloroform extraction to remove DNase activity. 1µg of RNA was then reverse transcribed into cDNA using SuperScriptIII reverse transcriptase (Invitrogen) and random hexamer primers. cDNA was quantified using primers to target genes (SNO2735-2736 for H2A.1, SNO3009-3010 for H2A.2), with relative levels normalized to that of the *ACT1* gene (SNO819-820).

### Chromosome loss assays

Chromosome stability was monitored in three ways. First, loss of the *ARG4* allele (on Chromosome 7) was determined by measuring the rate of loss of the ability of an *ARG4/arg*D heterozygous strain to grow on SD(-Arg) dropout medium. The *ARG4/arg*D heterozygous strains were created by integration of one copy *ARG4* from *C. dubliniensis* to the *arg*D*/arg*D locus of WT [2,2], [0,2], [2,0], *bub1,* and *sgo1* strains. *C.d.ARG4* gene addback fragments targeted to the *arg*D locus were generated by fusion PCR using primers SNO143-147 and pSN69 (*C.d.ARG4*) template, and correct insertion was verified by colony PCR using primers SNO143+SNO263 (5’) and SNO147+SNO264 (3’). The resultant Arg^+^ strains were streaked onto pre-spo medium and propagated at 37°C for 4 days. Cells were then re-suspended in sterile H_2_O, and plated for single colonies (∼200 per plate) on YEPD medium, with incubation at 30°C for 2 days. Colonies were replica-plated onto SD-ARG medium, and monitored for growth; colonies unable to grow on SD(-Arg) medium were scored as a positive event for chromosome loss.

The second method took advantage of the fact that *C. albicans* contains loci conferring sensitivity to the monosaccharide sorbose dispersed along Chromosome 5 ^38^; exposure to sorbose-containing medium selects for cells that have spontaneously lost one copy of this chromosome. *C. albicans* WT and mutant strains were grown on pre-spo for 4 days at 37°C, then resuspended in sterile H_2_O. 10^5^ or 10^6^ cells were plated on sorbose-containing medium, and a dilution was plated on YEPD medium to verify the total number of plated cells. Colonies on YEPD were counted after 2 days at 30°C, and colonies on sorbose-containing plates were counted after 10 days at 37°C. Chromosome 5 loss frequencies were calculated as the ratio of colonies observed on sorbose divided by the number on YEPD.

The third method was based on a previously described technique to visualize specific chromosomes using a chromosomally integrated TetO-tandem array and a TetR-GFP fusion protein^39^. In this case, we used a strain (QMY85) in which ∼120 tandem copies of TetO were integrated into the *his1*Δ locus of *C. albicans* reference strain SN87, under selection for the linked *C.d. HIS1* gene. This strain also contains a codon-optimized TetR-YFP fusion, expressed using a *SNU114* promoter and *ADH1* terminator, under selection for the linked *C. maltosa LEU2* gene. Association of the TetR-YFP fusion with the TetO array results in the appearance of one yellow fluorescent dot in yeast nuclei. A deletion mutation in the *H2A.2* ORF was then introduced into this strain using pSN383 as described above. Samples were analyzed by microscopy as described in the Fluorescence Microscopy section.

### Phylogenetic analyses

Sequences encoding H2A predicted amino acid sequences were individually obtained from NCBI (https://www.ncbi.nlm.nih.gov/), and aligned using a MUSCLE multiple sequence alignment ^40^. Multiple sequence alignments were visualized using the ESPRINT program ^41^. The phylogenetic tree was generated using a neighbor joining method, and visualized using the interactive tree of life online software ^42^. The distance measure represents substitutions per amino acid.

### Western Blot

Cell numbers were normalized based on OD_600_, and lysed by addition of HU buffer [8M urea, 5% SDS, 200 mM Tris-HCl, ph 6.8, 1.5% DTT], followed by boiling for 20 minutes. Samples were separated on 12.5% acrylamide gels at 120 V for 30 minutes and blotted onto nitrocellulose membranes (Thermo Scientific). Blotting was performed using a BioRad apparatus in Electroblot Buffer [27.5 mM Tris-Base, 192 mM glycine, 20% methanol] at 20 volts for 15 minutes. A primary mouse anti-FLAG antibody (Santa Cruz Biotech; 1:500 dilution) and a fluorescently labelled anti-mouse secondary antibody (LI-COR; IRDye800CW Goat anti-Mouse IgG) were used to detect H2A.2-FLAG. The primary antibody to H3 was a Rabbit anti-H3 (abcam; 1:5000 dilution), and was detected using a fluorescently labelled anti-goat secondary antibody (LI-COR; IRDye680 Goat anti-Rabbit IgG). Images were acquired using an Odessy CLx Imaging system (LI-COR).

### Ploidy analyses by PCR

Verification of *H2A.1, H2A.2, H2B.1,* and *H2B.2* gene dosage was performed by qPCR using the following primers: SNO2735-2736 for *H2A.1*, SNO3009-3010 for *H2A.2*, SNO2725-2726 for *H2B.1*, SNO2729-2730 for *H2B.2*, with levels normalized to that of *ACT1* (SNO819-820), which is located separately on chromosome 5. DNA was purified by phenol-chloroform extraction, and 8 ng was used as a template for qPCR reactions.

For whole genome ploidy analyses, an established primer set ^43^ was used, with primer pairs targeting either the left or right arm of each chromosome. For all chromosomes, the primers targeting the left arm of each chromosome were used, with the exception of chromosome 3, where both the left and right arm sets were used to confirm whole chromosome duplication. Using these primers, qPCR was performed using 8 ng of DNA template per reaction, and ploidy values were normalized to DNA from the strain SN152, a confirmed diploid control.

### Fluorescence Microscopy

*C. albicans* was grown in 30 ml cultures at 30°C with shaking at 200 r.p.m. for 4-5 hours in YEPD to OD_600_ of ∼1.0. Cells were then harvested by centrifugation, washed two times with PBS, spotted onto poly-L-lysine coated coverslips, and allowed to dry. Cells were then fixed with 4% paraformaldehyde (PFA) for 15 minutes, and washed two times with PBS. Vectashield (Vector Laboratories) was applied, followed by application and sealing of a cover slip. Images were acquired under 100X oil immersion objective using a Nikon TiE inverted microscope with Tokogawa spinning disk CSU-X1 using detection settings for YFP (excitation 514 nm, emission 527 nm) and DAPI. All images were processed with the ImageJ software (National Institute of Health); the presence of chromosomal YFP spots was counted manually, and >900 cells were evaluated per strain. Only un-budded cells were included in these counts.

### Propidium iodide staining and flow cytometry

Propidium iodide staining and flow cytometry were performed as previously described^44^ with some modifications. Briefly, mid-log phase cells (OD ∼2.0) were collected and washed two times with 50:50 TE (50 mM Tris, pH 8: 50 mM EDTA). Cells were pelleted and then fixed in 95% ethanol for 4 hours. Following two washes with 50:50 TE, cells were treated with 1 mg/ml RNase A for 3 hours, and then 1 mg/ml proteinase K for 15 minutes. Following two washes with 50:50 TE, cells were re-suspended in 50 mg/ml propidium iodide, and incubated overnight at 4°C. Stained cells (200 ml) were diluted in 500 ml of 50:50 TE and analyzed using a LSR/Fortessa/X-20. Ploidy values were estimated by comparing the ratio of peak locations in experimental samples to that of the diploid control SN152.

### Nocodazole and Fluconazole resistance assays

WT [2,2] and the strains [4,0] diploid, [0,4] diploid, *bub1*, and *sgo1,* [2,0], [2,0] + *H2A.2*, and [2,0] + *H2A.2*^S121A^ were diluted from overnight cultures, and grown to log phase (OD∼1.0) in a 30° C shaker at 200 r.p.m. in YEPD. Cells were then serially diluted onto YEPD, YEPD + 50 µM nocodazole, or 10 µg/ml fluconazole, and grown from 24 hours at 30° C. Plates were imaged using a Canon EOS Revel T5i camera, and image manipulations were performed using the ImageJ software (NIH).

### Statistical analyses

Except for ChIP-seq analysis described below below, p-values were generated by performing unpaired t-tests comparing strains with the relevant comparators. A p-value of 0.05 was used as a cutoff for significance.

### ChIP-Seq

ChIP was conducted as previously described^45^ with minor alterations. 40mL of Protein G DYNA beads (10002D, Thermo Fisher Scientific) were used in place of Sepharose beads for immunoprecipitation and washes were conducted on a magnetic stand. Post-Proteinase K treatment, samples were cleaned using Macherey-Nagel PCR clean up columns (740609.250S, Macherey-Nagel), eluted in 36mL of water, and libraries were made from these samples.

Immunoprecipitations were conducted with 10 μg anti-FLAG M2 antibody (F3165, Sigma), 10 μg anti-Histone H3 antibody (ab1791, Abcam), or 4 μg of anti-Cse4 (Laura Burrack). For all ChIP-Seq experiments, cells were propagated in YEPD overnight and seeded the next day at an OD600=0.1 in YEPD, pre-spo media, or YEPD + fluconazole (10μg/ml). Samples were propagated to an OD_600_=1.0 prior to harvesting.

Sequencing library preparation was conducted as previously described^46^. Library construction was performed in 2 replicates for each ChIP sample and one whole cell extract for each strain utilized. ChIP-Seq samples were normalized to their whole cell extracts and viewed on the IGV genome browser.

### ChIP-seq alignment

ChIP-seq data from a HiSeq 4000 were aligned to the *C. albicans* genome using Bowtie [1] with the following settings: *bowtie -p{0} -v2 -M1 --best --un {1}_multi_un.fastq --max {1}_multi.fastq {2} -q ^46^ --sam {1}_multi.sam* Conversion to BAM files, indexing and sorting were performed with SAMTools [2].

### Generation of bedgraph files

Bedgraphs were created using BEDTools [3]. Bedgraph files were first expanded to include each position in the genome and then smoothed using a rolling mean of 500 base-pairs using the python package pandas (rolling.mean function). Finally, each position in the bedgraph was normalized to the same position in the smoothed WCE bedgraph file to visualize fold enrichment.

### Whole genome tile analysis

**1**. *C. albicans* encodes two alleles of Histone H2A that differ in C-terminal amino acid sequence. Schematic of Bub1-H2A.1-Sgo1 interactions in other eukaryotes. *C. albicans* retains orthologs of both Bub1 (orf19.2678) and Sgo1 (orf19.3550). B) *C. albicans* histone genes are distributed over three chromosomes. *H2A.1* (orf19.1051), *H2B.1* (orf19.1052), *H4.1* (orf19.1061), and *H3.1* (orf19.1059) occur on Chromosome 1, *CSE4* (orf19.6163), *H2A.Z* (orf19.327), *H2B.2* (orf19.6924), *H2A.2* (orf19.6925), and *H3.A* (orf19.6791) on Chromosome 3, and *H4.2* (orf19.1854) and *H3.2* (orf19.1853) are on Chromosome R, which also contains rRNA genes. C*) C. albicans, C. dubliniensis*, and *C. tropicalis* evolved divergent alleles of H2A. Among these three yeasts, *H2A.1* (orange shading) and *H2A.2* (blue shading) are more similar to their orthologs in the two other species than to the H2A paralog in their own species. Scale bar designates substitutions per amino acid. D) Amino acid differences between the H2A paralogs of *C. albicans, C. dubliniensis,* and *C. tropicalis* (not shown) localize to the C-terminus and include replacement of the highly conserved serine/threonine 121 residue with lysine (asterisk). E) Deletion of *H2A.2* but not *H2A.1* produces a colony sectoring phenotype. Wild-type and homozygous deletion mutants were propagated on YEPD medium and incubated at 30°C.

**Figure 2**. A *C. albicans* mutant lacking *H2A.2* phenocopies spindle checkpoint mutants. A and Deletion of *H2A.2, BUB1, SGO1*, and replacement of *H2A.2* with a *H2A.2* S121A mutation result in similar colony sectoring phenotypes. Colony morphology (A) and % colony sectoring (B) are shown for wild-type *C. albicans*, the [4,0] strain that lacks *H2A.2*, the [0,4] strain that lacks *H2A.1*, knockouts of the spindle checkpoint genes, *BUB1* and *SGO1*, and the [2,0], and [2,0]+*H2A.2*S121A mutants. C, D, E, and F) Mutants lacking *H2A.2, BUB1*, and *SGO1* display increased aneuploidy. C) Percent of colonies that have lost the ability to grow on arginine-deficient medium, relative to growth on YEPD. Strains were plated on arginine-deficient medium or YEPD after 4 days on pre-sporulation medium. Because these *ARG4/*D strains contain the *ARG4* gene on only one copy of chromosome 7, loss of that chromosome renders cells auxotrophic for arginine. D) Percent of colonies that grow on sorbose-containing medium, relative to growth on YEPD. Because cells with both copies of chromosome 5 are killed by sorbose, this medium selects for cells that have lost one copy of the chromosome. E) Top, Superimposed fluorescence and phase microscopy of QMY85, which contains tandem copies of the tet operator on Chromosome 5, as well as the tet repressor fused to YFP. A yellow spot marks one of two copies of Chromosome 5 (white arrow) and the nucleus is stained with DAPI (blue arrow). Bottom, example of neighboring cells with 1 or 2 spots. F) % of cells with two spots, counting un-budded cells only. G) Sensitivity to the microtubule poison Nocodazole (50 µM) shown by serial dilution.

**Figure 3**. H2A.1 and H2A.2 play opposing roles in the chromosome stability of tetraploid cells and in the acquisition of resistance to fluconazole. A) Cartoon of the *C. albicans* parasexual cycle, which consists of mating between diploid **a** and a cells to form tetraploid **a**/a cells, followed by ploidy reduction by concerted chromosome loss rather than meiosis, as occurs in other eukaryotes. B) Viability of WT, tetraploid, or tetraploid cells containing only H2A.1 ([8,0] Tetraploid), or H2A.2 ([0,8] Tetraploid) on YEPD (left) vs. pre-sporulation medium (right). Plated cells were propagated for four days at 37°C, and survival was calculated as the ratio of colonies on each medium to the estimated number of plated cells determined using a hemocytometer. C) Chromosome stability of *ARG4/arg4*D WT, tetraploid, and derivative strains on pre-sporulation medium after four days, as measured by growth on SC(-Arg) medium, indicative of the loss of one copy of chromosome 7. D) Serial dilutions of *C. albicans* WT, [4,0] diploid, [0,4] diploid, *bub1, sgo1*, [2,0] diploid, and [2,1] diploid strains on YEPD or YEPD + fluconazole (10 µg/ml).

**Figure 4.** Impact of ploidy and environment on histone expression and deposition. A) RT-qPCR analysis of expression of *H2A.1* and *H2A.2* in the indicated genotypes and conditions. “D” represents a wild-type [2,2] Diploid strain and “T” a wild-type [4,4] Tetraploid strain. The average results for three biological replicates is presented after normalization to H3 mRNA, along with the standard deviation. R.u. stands for relative units. B) Immunoblot of levels of H2A.2-FLAG in the indicated genotypes and conditions. Quantification using a LI-COR imager is shown on the right. C) ChIP-seq analysis. Plots of bedgraphs of normalized read densities are shown for the indicated genotypes and conditions. Data represent the results of one replicate of each sample for *CEN5*. Data for all centromeres are shown in Figure S6, and scatterplots of replicate data are shown in Figure S7. D) Boxplots showing normalized ChIP-seq enrichments (relative to the whole-cell extract sample, see Materials and Methods) of nonoverlapping 5 kb tiles of the *C. albicans* genome separated into those overlapping with annotated centromeres vs. the remainder of the genome.

